# Transcriptomic contributions to a modern cytoarchitectonic parcellation of the human cerebral cortex

**DOI:** 10.1101/2023.03.22.533710

**Authors:** Leana King, Kevin S. Weiner

## Abstract

Transcriptomic contributions to the anatomical, functional, and network layout of the human cerebral cortex (HCC) has become a major interest in cognitive and systems neuroscience. Here, we tested if transcriptomic differences support a modern, algorithmic cytoarchitectonic parcellation of HCC. Using a data-driven approach, we identified a sparse subset of genes that differentially contributed to the cytoarchitectonic parcellation of HCC. A novel metric (cortical thickness/myelination ratio; CT/M ratio), as well as cell density, correlated with gene expression. Enrichment analyses showed that genes specific to the cytoarchitectonic parcellation of the HCC were related to molecular functions such as transmembrane transport and ion channel activity. Together, the novel relationship between transcriptomics and the CT/M ratio bridges the gap among i) gradients at the macroscale, ii) areas at the meso-scale, and iii) cell density at the microscale, as well as supports the recently proposed cortical spectrum theory.

## Introduction

Transcriptomic contributions to the anatomical, functional, and network layout of the human cerebral cortex (HCC) has become a major interest in both cognitive and systems neuroscience in the last decade (Fornito et al., 2019; Arnatkeviciute et al., 2021c; Deco et al., 2021; Wagstyl et al., 2022). Recent research shows that active transcription of a small set of genes contributes to large-scale gradients and functional hierarchies across the HCC, which likely contribute to the regional functional and structural differences in the HCC. Specifically, previous findings show that a subset of genes increase or decrease their expression from primary to association cortices (Burt et al., 2018), likely also contributing to the hierarchical organization of functional processing streams (Gomez et al., 2019). Nevertheless, it is largely unknown if transcriptional differences also contribute to modern, algorithmic definitions of cytoarchitectonic areas across the whole HCC (Amunts et al., 2020).

This gap in knowledge persists for at least three main reasons. First, advancements in sequencing methodologies were only developed recently and freely shared with the larger neuroscience and human brain mapping fields, which enables novel links between gene expression in human postmortem tissue to current in-vivo measures of cortical anatomy and function (Hawrylycz et al., 2012; Bludau et al., 2018; Arnatkeviciute et al., 2019; Wang et al., 2022). Second, most human research that has linked transcriptomics to brain-wide features have focused on either connectivity (Arnatkeviciute et al., 2021b; Oldham et al., 2022), functional differentiation (Burt et al., 2018), or a small subset of cortical areas (Zachlod et al., 2022). Third, a modern, observer-independent cytoarchitectonic parcellation of the HCC was only recently published in the last two years. For example, classic cytoarchitectonic approaches (Brodmann, 1909; von Economo and Koskinas, 1925) rely on the eyes of the anatomists to qualitatively determine when two adjacent pieces of cortex are cytoarchitectonically distinct. These classic maps still used in cognitive and systems neuroscience have additional flaws aside from their qualitative nature. For example, while 60-70% of HCC is buried in indentations, or sulci (Welker, 1990; Armstrong et al., 1995), Brodmann did not even examine sulci, despite the fact that thousands of studies continue to cite his map (Abbott, 2003; Zilles and Amunts, 2010; Zilles, 2018). Contrary to these classic approaches, recent methods use observer-independent algorithms to statistically determine a cytoarchitectonic boundary between adjacent pieces of cortex. Importantly, the modern cytoarchitectonic parcellation of the HCC is functionally meaningful at both the areal and network level. Determining if transcriptomic differences contribute to a modern cytoarchitectonic parcellation of the HCC can guide the broad fields of cognitive and systems neuroscience in understanding the complex relationship among gene expression, cytoarchitecture, and functional networks of the HCC, in addition to targeting genes that could be impacted in deficits altering this multimodal and multiscale relationship (Seidlitz et al., 2020; Arnatkeviciute et al., 2021a).

To address this gap in knowledge, we integrated transcriptional data and cytoarchitectonic parcellations of the HCC from two freely available datasets (Hawrylycz et al., 2012; Amunts et al., 2020). Using a data-driven approach, we identified a sparse subset of genes that differentially contributed to the cytoarchitectonic parcellation of the HCC. These genes were located in two distinct clusters that formed opposing gradients of expression. Interestingly, these gradients were also correlated with a metric that considered the ratio between cortical thickness and myelination. Clustering analyses of cytoarchitectonic areas revealed both within-network and between-network relationships. When repeating our analyses with a recent multimodal parcellation of the HCC (Glasser et al., 2016), ∼90% of genes were shared between both parcellations, while ∼10% were specific to each parcellation.. Enrichment analyses showed that genes specific to the cytoarchitectonic parcellation of the HCC were related to molecular functions such as transmembrane transport activity and ion channel activity, while genes specific to the multimodal parcellation of the HCC were related to calcium channel activity and cellular components such as neuron development. Altogether, our results identify a relationship between genetic expression and an observer-independent cytoarchitectonic parcellation of the HCC for the first time (to our knowledge), as well as identify novel genes for future studies interested in further understanding the contribution of genetic expression to different anatomical, functional, and multimodal parcellations of the HCC.

## Results

### A sparse subset of brain-specific genes contributes to a modern cytoarchitectonic parcellation of the human cerebral cortex (HCC)

To identify genes that potentially contribute to an algorithmic, observer-independent cytoarchitectonic parcellation of the HCC (Figure 1*A*), we assessed gene expression profiles that were significantly different among cytoarchitectonic areas (cROIs) using two freely available datasets: 1) a modern cytoarchitectonic parcellation of the HCC from 10 post-mortem brains (JüBrain: https://www.fz-juelich.de/inm/inm-1/EN/Home/home_node.html) and 2) the Allen Human Brain Atlas, which consists of transcriptomic data from tissue samples in 6 post-mortem brains (AHBA: http://brain-map.org/). To test if there is a relationship between genetic expression and the observer-independent cytoarchitectonic parcellation of the HCC, transcriptomic data were pre-processed and aligned to the cytoarchitectonic parcellation (Figure 1*B*) via the abagen package. This process precisely aligned the expression of 15,630 genes (controlling for false positives, duplicate probes, etc; Materials and Methods) across 111 cROIs.

**Figure 1.**
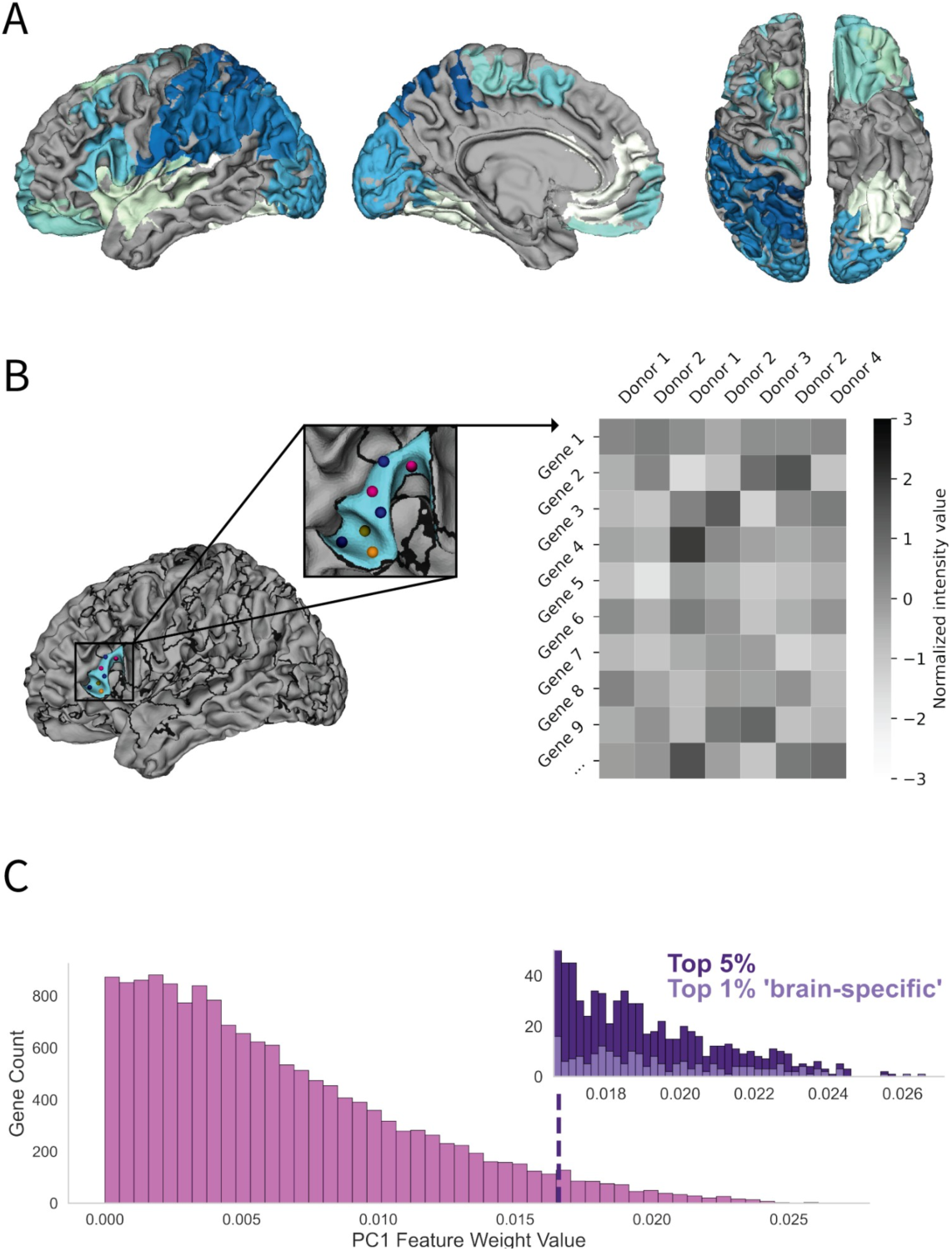
Gene selection within cytoarchitectonic neighborhoods of the human cerebral cortex. (A) A recent, observer-independent approach (Amunts et al., 2020) parcellated the human cerebral cortex (HCC) into 111 cytoarchitectonic areas (Materials and Methods). (B) *Left:* Each tissue sample from 6 post-mortem brains was aligned to this novel cytoarchitectonic parcellation (Materials and Methods). Example locations of several different probes from different donors are depicted within area 45 (“Broca’s” area) within the inferior frontal cortex. Each color corresponds to a single donor, with each probe containing the relative intensity value of the microarray expression for the genes surveyed. *Right:* Matrix illustrating the transcriptomic data across the different probes within a single region. Each column corresponds to one of the probes shown on the left. Each row is the normalized expression value of a single gene. (C) A histogram of the feature weights for PC1 are shown (pink) along with the selection of the top 5% of genes with the highest feature weight values (dark purple). The top 200 genes that were most differentially expressed across all 111 cROIs were selected and used for the remainder of the analyses after filtering for ‘brain-specific’ genes (Burt et al., 2018).

A Principal Components Analysis (PCA) was implemented to identify genes that were most differentially expressed across all 111 cROIs. A histogram (Figure 1 *C*) of resultant feature weights for each gene empirically supported that a majority of the genes do not contribute to the primary axis of transcriptomic variation. In order to identify genes that most strongly contributed to the cytoarchitectonic arealization of HCC, we restricted our analyses to the top 1% (n=200) of brain-specific (as identified by Burt et al., 2018) genes that were most differentially expressed as in our previous work (Gomez et al., 2019, 2021) (Figure 1*C*, light purple). Importantly, variation in expression of the identified genes across the geodesic surface were not driven by trends in spatial autocorrelation (Supplementary Figure 3*B*).

### Observer-independent cytoarchitectonic areas are embedded within two opposing transcriptomic gradients

As previous research showed that cROIs in the visual processing hierarchy are located within opposing transcriptomic gradients (Gomez et al., 2019), we tested the targeted hypothesis that this relationship extended more broadly to cROIs distributed throughout HCC. To test this hypothesis, the top 200 genes were submitted to an agglomerative hierarchical clustering algorithm (Materials and Methods). This approach revealed that genes clustered into two groups at the highest level with branch distances much greater than that of other clustering levels (Figure 2*A*). These two clusters generated two opposing gradients among cROIs: one that increased in expression and another that decreased in expression along the order determined by score values from PC1 (Figure 2*B*). Interestingly, however, only 20% of genes overlapped between the present group of genes and those identified previously that contributed to the positioning of areas in the visual processing hierarchy (Gomez et al., 2019). We also tested the correlation between the hypothesis matrix and the observed matrix and found an R-value of 0.87 with a range of 0.74 to 0.92 (Figure 2*C*) when randomly selecting for 50% of the cROIs (permuted N = 1,000). Furthermore, we generated a null distribution of correlations between the hypothesis matrix and shuffled cROIs, which was well below the range observed for 50% of the ordered cROIs, indicating the robustness of the identified relationship. For comparison, we also applied the same clustering method, but to the bottom 200 genes, which revealed a much more homogenous and smaller branching distance across levels (Figure 2*A*), further supporting that this relationship was specific to the genes most differentially expressed among cROIs distributed throughout the HCC.

**Figure 2.**
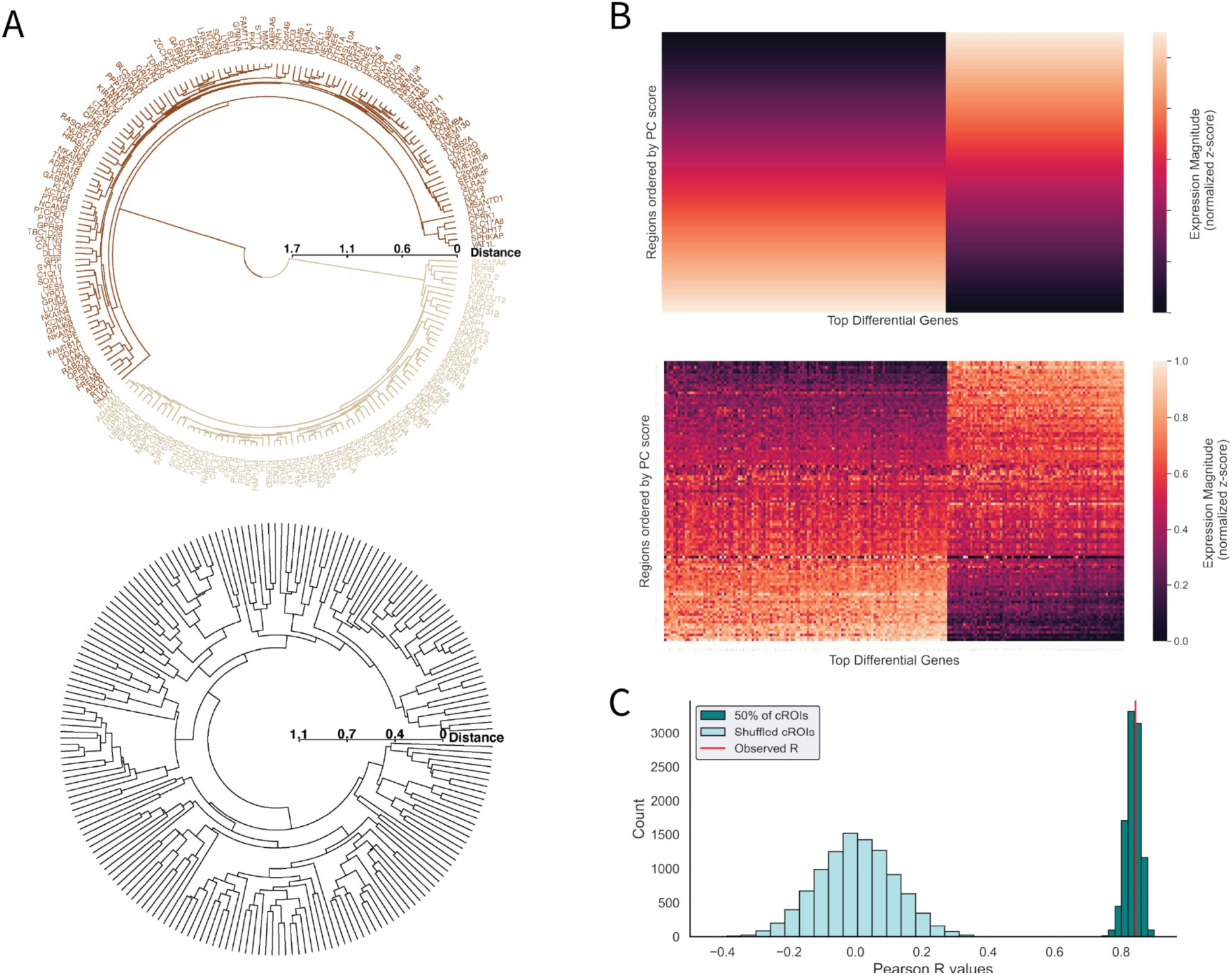
Binary clustering of the top 200 genes reveals that opposing expression gradients contribute to the cytoarchitectonic layout of cortical areas identified using an observer-independent technique. (A) Dendrograms showing the algorithmic clustering of the top and bottom 200 genes. *Top*: The top 200 genes that are most differentially expressed cluster predominantly into two groups at the highest level (brown and tan). The names of each gene are plotted radially. *Bottom:* On the contrary, hierarchical clustering reveals minimal distance between branches for the bottom 200 genes, indicating very little transcriptomic differences between clusters in the dendrogram. (B) *Top:* Hypothesis matrix of genetic gradients (based on Gomez et al., 2019). The hypothesis matrix was created using a stepwise change in gene expression for each cROI as ordered by their PC score in which one set increases, and the other set decreases, in expression across regions. *Bottom:* Observed matrix of the gene expression of the top 200 differentially expressed genes (y-axis) across the 111 cROIs ordered by their score from PC1 (x-axis). The brown cluster of genes from (A) represents the increasing cluster on the left, while the tan cluster from (A) represents the descending cluster on the right. (C) We calculated the correlation between the hypothesis matrix and the observed matrix by randomly selecting from 50% of the cROIs to correlate with corresponding rows in the hypothesis matrix and bootstrapped with an N=1,000. This resulted in a distribution of values ranging from 0.74 to 0.90 with an observed R value of 0.84 (p < 0.0001). We also generated a null distribution (N = 1,000; cyan) of R values by shuffling the cROI order in the observed matrix prior to correlating it to the hypothesis matrix.

### Primary axis of transcriptomic variation reveals within and between network clustering of cytoarchitectonic areas

PCA on the cROIs with each gene set as a feature identified that the first PC explained 24.12% of the variance. As such, we refer to PC1 as a representation of the primary axis of transcriptomic variation in the cytoarchitectonic arealization of HCC. There is a clear structure among the distribution of cROIs along this primary axis: early visual areas were positioned on one end of the transcriptomic axis (Figure 3*A*, hOc1 (e.g, striate cortex, or V1), hOc2 (V2), etc., lower right column), while frontal and insular regions were positioned on the other end of the axis (Figure 3*A*, Ia2, Op7, etc; upper right column).

**Figure 3.**
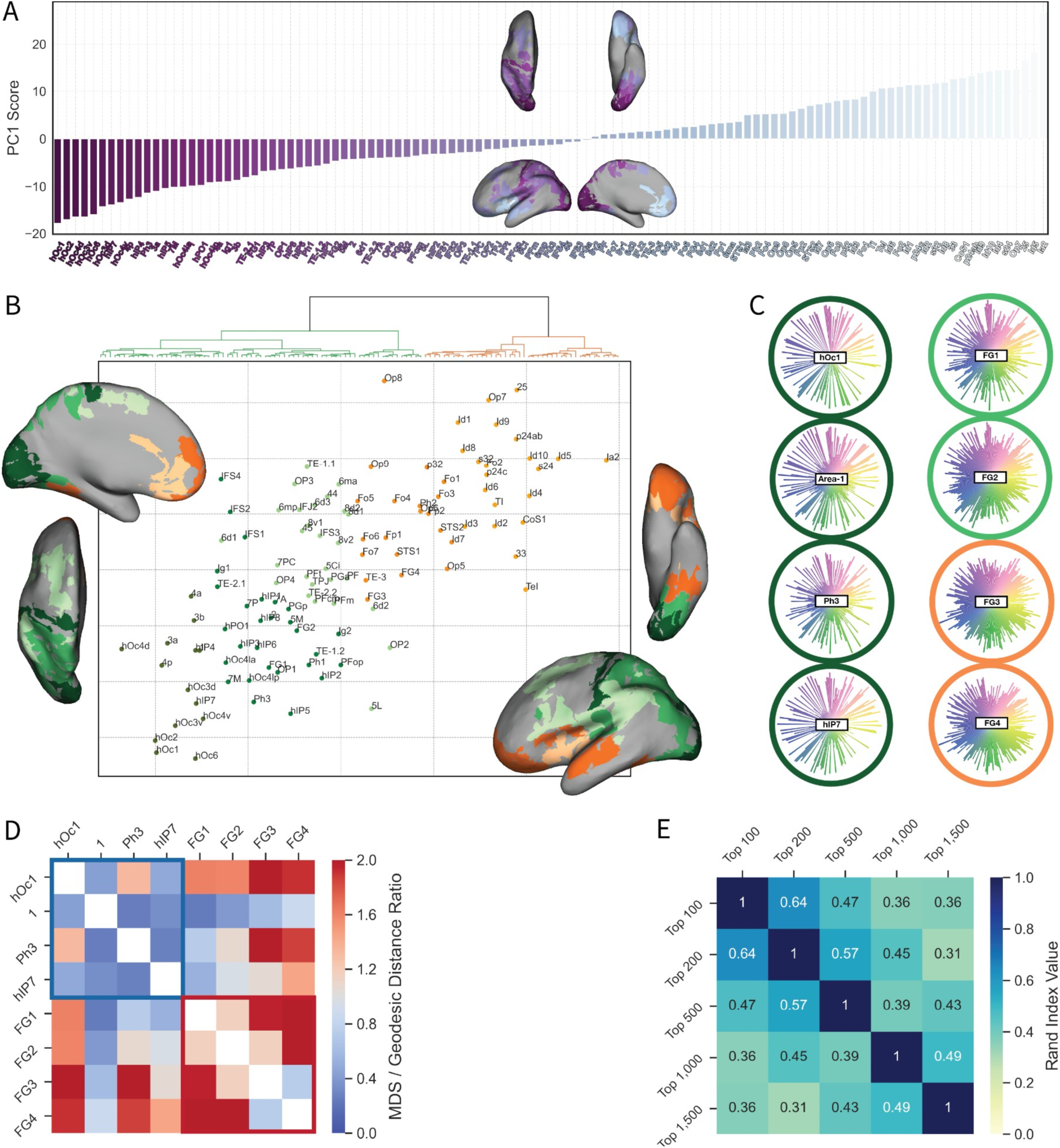
Hierarchical clustering of transcriptomic fingerprints reveals within and between network clustering of cytoarchitectonic areas. (A) Scores obtained from the first PC, highlighting the linear order of cROIs that is used for visualizations in figures throughout the manuscript. Score values from the first PC (y-axis) are plotted as a function of each cROI (x-axis), ordered by score values (from lowest (left) to highest (right)). Names of the cROIs ordered by the first PC are illustrated on the bottom axis starting with hOc1 on the left in dark purple and ending with insular region Ia2 in light blue on the far right. Also shown are topographic plots that illustrate the score value for the cROIs on the MNI152 inflated cortical surface. cROIs in dark purple have a lower PC score while cROIs in light purple have a higher PC score. (B) Agglomerative hierarchical clustering was applied to the cROIs and their expression of the 200 most differentially expressed genes, which revealed two main clusters (orange and green) at the highest level (top) with subclusters shown in different shades of green and orange. Topographical representations of the cROIs with their corresponding cluster label are also shown in different views. To further illustrate the genetic differences and similarities between cROIs, multidimensional scaling was applied to cROIs and their differential gene expression and then color-coded using labels from the hierarchical clustering. Regions close together in the 2-D plane are more transcriptomically similar and regions further apart are more transcriptomically dissimilar. Within the MDS, there are two main types of genetic similarities: within-network and between-network. For example, all the early visual regions cluster together in the dark green cluster on the left highlighting within-network similarities. Additionally, while regions FG1, FG2, FG3 and FG4 are cortically adjacent to one another, FG1 and FG2 are in one transcriptomic cluster (green), while FG3 and FG4 are in a transcriptomically distinct cluster (orange). The former areas cluster more closely to parietal areas, while the latter cluster more closely with areas in superior temporal cortex, both of which are indicative of between network clustering. (C) Transcriptomic fingerprints of selected cROIs in which the radial axis shows the relative gene expression for the top 200 differential genes. The column on the left highlights regions within the same cluster, while the column on the right highlights regions across two different clusters to qualitatively illustrate the similarities and differences in gene expression profiles. (D) Ratio comparison between geodesic distance of cROIs and their transcriptomic similarity as measured by the Euclidean distance between the cROIs on the MDS 2-D plane. Values in red (or above one) have a greater transcriptomic difference than geodesic distance (i.e. across network clustering) while values in blue (or less than one) have a greater geodesic distance than transcriptomic difference (i.e. within network clustering). (E) Rand index matrix representing the pairwise stability of cluster labels when selecting for a different number of the highest feature weight values (N = 100, 200, 500, 1,000 and 1,500 genes), while still limiting analyses to brain-specific genes (N =1,898).

To further explore the relationship of this primary axis of transcriptomic variation to the arealization of the HCC, we generated transcriptomic profiles - or *fingerprints* (Figure 3*C*) - for each cROI consisting of the normalized average expression of the top 200 differentially expressed (DE) genes. Using these transcriptomic fingerprints, we then performed an agglomerative hierarchical clustering algorithm to examine the different clusters of cROIs based on the similarity of transcriptomic fingerprints. These similarities were further represented in a 2-dimensional plane using Multi-Dimensional Scaling (MDS) in which the Euclidean distance between regions represented the transcriptomic similarity or dissimilarity to other regions.

This analysis revealed two primary clusters of cROIs based on transcriptomic fingerprints. One cluster represented in orange in Figure 3*B* primarily consists of regions from frontal, cingulate, and insular regions, while the other cluster represented in green contains regions from occipital, parietal, and temporal regions. Furthermore, within each cluster, there are smaller subdivisions (various shades of green and orange in Figure 3*B*). When further exploring the clustering within the MDS, we found that there were two types of genetic similarities. One shows ‘within transcriptomic network’ clustering, such as visual regions that cluster together in the dark green cluster in the lower left of Figure 3*B*. The other shows ‘across transcriptomic network’ clustering such as fusiform regions (FG1, FG2, FG3, and FG4). For example, even though both areas FG2 and FG4 are located in the lateral FG, contain regions that are selective for faces and words, and are cortically adjacent to each other (Weiner et al., 2017), they are located within different transcriptomic networks. These qualitative differences can be observed in the transcriptomic fingerprints highlighted in Figure 3*C*.

To test if spatial proximity contributed to transcriptomic similarity, we calculated the ratio between the pairwise Euclidean distance between two cROIs in the MDS plane and the corresponding geodesic distance on the surface (Figure 3*D*). Both distance metrics were normalized. Values in red (greater than 1) correspond to a greater transcriptomic difference over geodesic distance while values less than 1 (in blue) correspond to a greater geodesic distance than transcriptomic difference. Pairwise ratio values are shown for the 8 representative cROIs that were also illustrated in Figure 3*C*. Ultimately, while there are a few cROIs that are as transcriptomically similar as they are proximal, for a majority of regions, geodesic distance does not determine transcriptomic similarity. For example, regions hOc1 and Area 1 are cortically far apart, but transcriptomically similar; however, regions FG1 and FG2 are cortically proximal to FG3 and FG4, but are transcripitomically distinct from each another.

For cROI ‘across network clustering’ pairs, we quantified the relationship between transcriptomic similarity and cortical distance by calculating a transformed ratio value (absolute value of the Pearson’s correlation coefficient in gene expression between two regions divided by the normalized geodesic distance along the surface) and then subtracted this value from 1 (Supplementary Figure 2*A*). 0 represents region pairs that are both highly correlated and cortically distant. The region pair to exhibit the strongest ‘across network clustering’ are regions 4p of the somatosensory cortex and Ph3 of the parahippocampal cortex (R = 0.81 and normalized geodesic distance = 0.77) that cluster together with early visual regions and other somatosensory regions. Other surprising region pairs include STS2 in superior temporal cortex with p32 in medial anterior cingulate cortex, hOc3v in extrastriate visual cortex with 4a in primary motor cortex, and hOc1 in striate cortex with 3b in primary somatosensory cortex. The combination of these findings indicate that transcriptomic similarity between region pairs is not driven by cortical proximity.

Lastly, we repeated our analyses with a variable number of genes (N = 100, 500, 1,000 and 1,500) to test if the number of genes affected the clustering of regions. Compared to our analyses with 200 genes, we found that the clusters remain stable across the different Ns observed with decreasing Rand Index values as the number of genes increased (Figure 3*E*). This finding further enhances the reliability of the clustering results.

### CT/M ratio: A novel metric that is highly correlated with the primary axis of transcriptomic variation

What underlying anatomical features are related to this primary axis of transcriptomic variation? While previous research has explored the relationship between transcriptomics and gradients of cortical thickness and myelination {Burt et al., 2018; others}, our results show that striate cortex and primary motor cortex are transcriptomically similar to one another. This is somewhat surprising since striate cortex is cortically thin, heavily myelinated, cell dense, and has a prominent and subdivided layer IV (Collins et al., 2010; Balaram and Kaas, 2014; Gomez et al., 2021), while primary motor cortex is cortically thick, heavily myelinated, relatively cell sparse (compared to striate cortex) and the presence and prominence of layer IV is contentious (García-Cabezas and Barbas, 2014; Barbas and García-Cabezas 2015). Given recent work showing that the ratio between cortical thickness and myelination (CT/M ratio) was a particularly useful metric for parcellating one HCC area from another (Willbrand et al., 2022), we hypothesized that despite these differences in anatomical and architectonic features across cortical areas, transcriptomic similarities may be capturing a ratio of anatomical features. To test this hypothesis, we aligned our cROIs to average cortical thickness and myelination maps from the 1,096 participants included in the Human Connectome Project (HCP). This analysis revealed a strong positive correlation between the expression of these 200 genes and the CT/M ratio across cROIs of the HCC as shown in Figure 4*A* (R=0.872; *p* < 0.01e^-34^).

**Figure 4.**
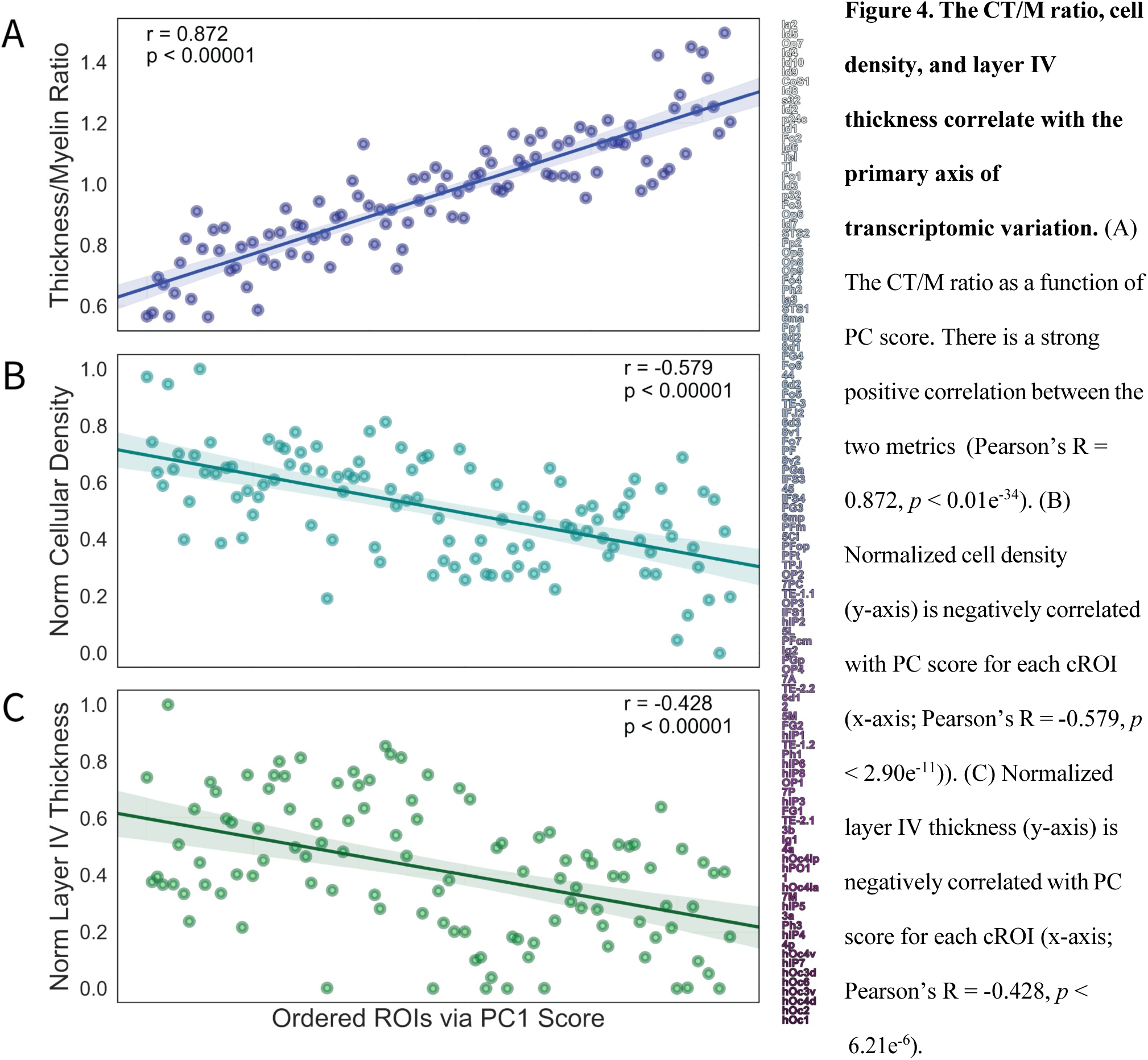
The CT/M ratio, cell density, and layer IV thickness correlate with the primary axis of transcriptomic variation. (A) The CT/M ratio as a function of PC score. There is a strong positive correlation between the two metrics (Pearson’s R = 0.872, *p* < 0.01e^-34^). (B) Normalized cell density (y-axis) is negatively correlated with PC score for each cROI (x-axis; Pearson’s R = -0.579, *p* < 2.90e^-11^)). (C) Normalized layer IV thickness (y-axis) is negatively correlated with PC score for each cROI (x-axis; Pearson’s R = -0.428, *p* < 6.21e^-6^).

In addition to this novel CT/M ratio, underlying cytoarchitectonic features likely contributing to this primary transcriptomic axis are cell density and the presence/prominence of layer IV. For example, striate cortex (lower left of Figure 4*A*) has a uniquely prominent layer IV with several sublayers, as well as is the most cell dense area in the HCC (Collins et al., 2010; Balaram and Kaas, 2014; Gomez et al., 2021). Additionally, visual areas have a higher cell density than HCC areas outside of visual cortex (Collins et al., 2010). Further, area Ia2 does not have a layer IV (agranular) while areas Id5 and OP7 have a thin layer IV (von Economo, 2009; Zilles and Amunts, 2012; Amunts and Zilles, 2015) (dysgranular; upper right of Figure 4 *A*). Thus, to further test if there was also a relationship between cell density, granularity, and the primary axis of transcriptomic variation of the cROIs, we incorporated cell density and layer IV thickness estimates from the classic Von Economo - Koskinas atlas into our computational, multimodal pipeline (Materials and Methods). This analysis showed a negative correlation (R = -0.579, *p* < 2.90e^-11^) between cell density and the primary axis of transcriptomic variation (Figure 4*B*). Layer IV thickness was also negatively correlated with the primary axis of transcriptomic variation (R = -0.428, *p* < 6.21e^-6^ ; Figure 4*C*). Altogether, the primary axis of transcriptomic variation among cROIs likely reflects aspects of cell density, the presence/prominence of layer IV, and the novel metric of the CT/M ratio.

### Gene enrichment analyses reveal that different sets of genes contribute to cytoarchitectonic and multimodal HCC parcellations

To gain a deeper understanding of the functional roles of the top 200 differentially expressed genes contributing to the observer-independent cytoarchitectonic parcellation of the HCC, gene symbols were submitted to an enrichment analysis resulting in five clusters of unique structural or functional properties (Materials and Methods). The most significant terms were associated with channel and synaptic function, cellular projections and maintenance, and transport of molecules or ions (Figure 5*A*).

**Figure 5.**
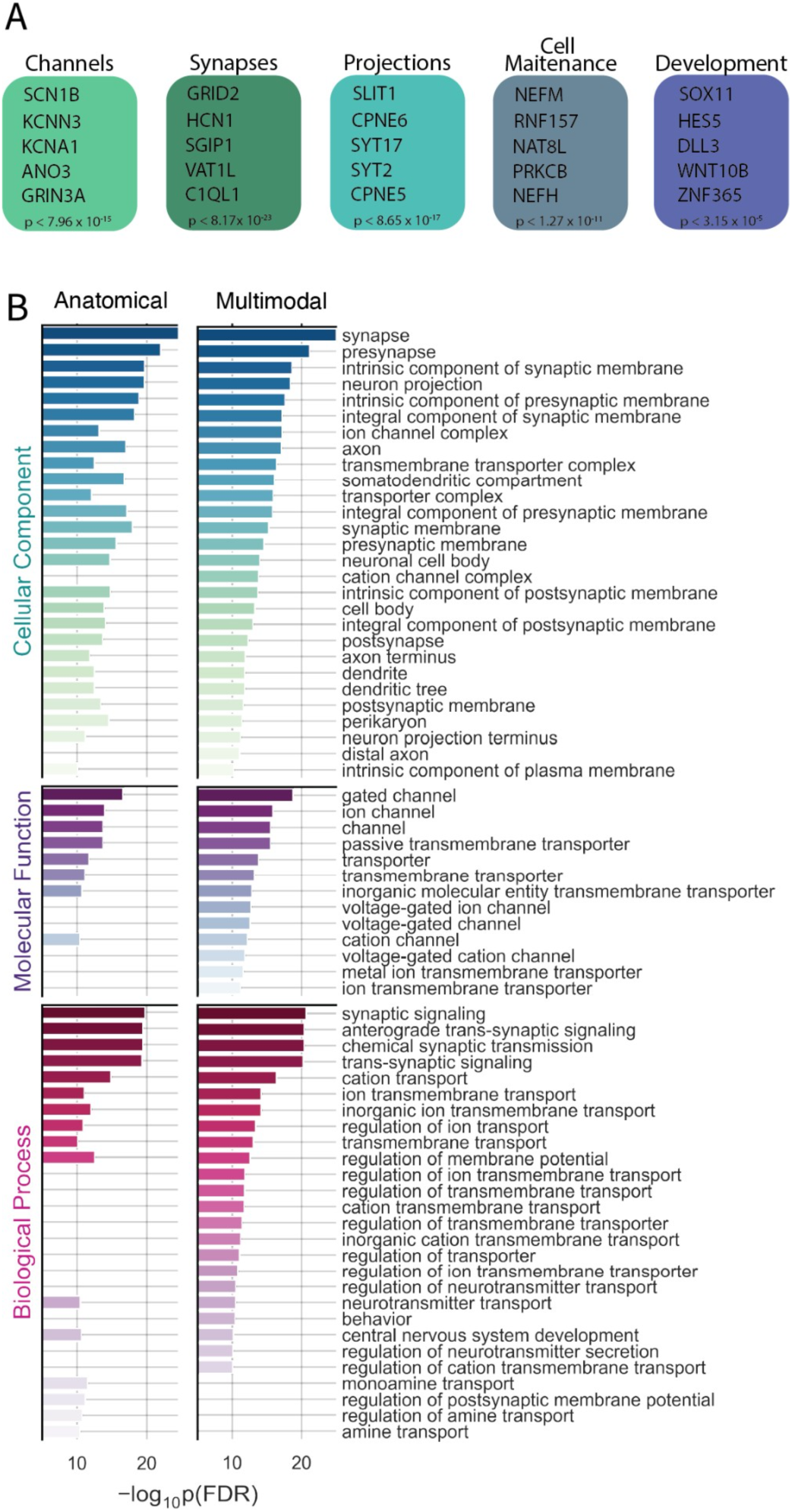
*Cytoarchitectonic* genes are more enriched in cell body functions and amine transport, while *multimodal genes* are more enriched in ion channel and transport functions. (A) List of 5 key annotation categories (along with example genes associated with each category from the Gene Ontology (GO) Database) for the 180 genes overlapping between the cytoarchitectonic parcellation and the multimodal parcellation (Supplemental Figure 5). Top categories include ion channel function, synaptic function, cellular projections, cell maintenance, and brain development. (B). Gene enrichment results for the differing sets of genes for the cytoarchitectonic (left column) and multimodal (right column) parcellation of the HCC. Each GO term is plotted by the log-transformed FDR-corrected p-value in the horizontal bars for their respective significance in either set of genes (i.e. cytoarchitectonic or multimodal). Given that 180 of the genes are the same between both sets, enrichment results are similar for each set in which both are heavily enriched in synaptic functions. Yet, differences between the two sets show that the ‘anatomical’ gene set is more enriched in cell body functions and amine transport while the ‘multimodal’ genes are more enriched in ion channel and transport functions.

To test if these genes were specific to the cytoarchitectonic parcellation of the HCC or generalized to additional parcellations of the HCC, we repeated our analyses with a multimodal parcellation of the HCC (Glasser et al., 2016; Supplemental Figure 4). Our quantifications revealed that roughly 90% of the top 200 genes are shared between parcellation modality, while roughly 10% are different. Specifically, we repeated the enrichment analyses with different sized sets of differential genes (N=100, N=200, N=500, N=1,000; Supplemental Figure 5). Through each iteration, at least 8% of genes are unique to either parcellation with the most being 14%. Interestingly, enrichment analyses showed that parcellation-specific genes had different functions: cytoarchitectonic arealization genes were enriched in molecular functions such as transmembrane transport activity and ion channel activity while the multimodal arealization genes were enriched in functions such as calcium channel activity and cellular components such as neuron development (Figure 5*B*). Furthermore, we submitted the top 200 differentially expressed genes of the cytoarchitectonic parcellation to a protein-protein network analysis via the STRING database (Supplemental Figure 6). We found that the most connected network of genes performed ion-channel functions and belonged to the KCNA and SCN family, while the second most connected network consisted of neurotransmitter-related genes followed by cell maintenance genes.

## Discussion

In the present study, we implemented a data-driven approach and identified transcriptomic contributions to a modern algorithmic cytoarchitectonic parcellation of the HCC. These findings provide two clear bridges to recent parallel empirical tracks examining the contributory role of transcriptomics to identified gradients and maps of the HCC. The first track explores the relationship between transcriptomics and gradients in the HCC (Burt et al., 2018; Gomez et al., 2019; Vogel et al., 2022). The second track explores the relationship between transcriptomic-based parcellations (TBP) of the HCC relative to modern and classic areal parcellations of the HCC (Shine et al., 2022; Gryglewski et al., 2018, 2022). The first bridge is that neither track considered a modern, observer-independent cytoarchitectonic parcellation of the HCC – instead, they considered observer-dependent parcellations that have several shortcomings as discussed previously (Amunts and Zilles, 2015; Amunts et al., 2020). The second bridge is that neither track (to our knowledge) considered a combinatory metric (the ratio between cortical thickness and myelination; CT/M ratio) integrating the smooth gradients and the discrete, parcellated areas. The two bridges provided by the present study integrate these two empirical tracks in a critical manner and provide theoretical insights for future studies. Below, we discuss these findings in the context of: i) the role of transcriptomics and the CT/M ratio for integrating continuous gradients and discrete, classic “crazy-pavement maps” along a modern “cortical spectrum,” ii) transcriptomics as a bridge connecting areas of functionally distinct cortical networks, and iii) transcriptomics contributing to different scales of organization and different parcellations of the HCC.

### The role of transcriptomics and the CT/M ratio for integrating continuous gradients and discrete, classic “crazy-pavement maps” along a modern “cortical spectrum”

In the broader neurology, cognitive neuroscience, and human brain mapping communities, the last decades continue a longstanding trend of identifying gradients and parcellations of the HCC. For example, in a review of Bailey and Bonin’s *Isocortex of Man*, Le Gros Clark (1952) wrote: “Finally, neurologists should continue to regard with the greatest suspicion the incredibly complicated “crazy-pavement” maps of cortical areas which have from time to time been elaborated by the Vogt school since Campbell’s and Brodmann’s original studies were first published” (LeGros Clark, 1952, pp. 104). Both Brodmann and Sanides (among others) were students of the “Vogt school” and along with the Vogts, produced a parcellation of the HCC into ∼180-200 areas over several decades (Nieuwenhuys, 2013; Foit et al., 2022) – a number that is comparable to maps produced with modern multimodal and data-driven methods (Glasser et al., 2016; Huth et al., 2016). Despite this “crazy-pavement” map, the Vogts (Vogt and Vogt, 1919) also acknowledged that the boundaries among adjacent cortical areas were not sharp and building on their hypothesis, Sanides (1962) proposed that in addition to cortical areas, there were directions of “gradation” streams that correlated with directions of brain evolution and neocortical development (summarized by Henssen et al., 2016).

A recent study (John et al., 2022) proposed the concept of a *cortical spectrum* that builds on the foundation of this classic work by integrating three features of functional and structural organization: i) a mosaic of functionally and structurally distinct parcellations of the cortical mantle, ii) a seemingly uniform six-layered structure in which connectivity is the driving factor of functional differences (Barbas, 1986; Barbas and Rempel-Clower, 1997), and iii) gradients of architectonic structure that predict cortico-cortical connections. Here, we propose that our results nicely complement and extend these classic and modern theoretical and empirical findings linking different scales of brain structure and function by incorporating the correlated relationship between gene expression and the CT/M ratio.

Specifically, the primary axis of transcriptomic variation as captured by PCA identified in the present study is strongly associated with the granularity, or lamination, of cROIs. For example, highly granular regions, such as hOc1 (primary visual cortex) and 4a (a subarea of primary motor cortex), appear on one end of the spectrum while more dysgranular or agranular regions, such as many insular and frontal cROIs in limbic cortices, appear on the other end. These results are consistent with the *cortical spectrum* proposal, as well as two recent transcriptomic findings in three main ways.

First, John and colleagues (2022) implemented a data driven approach and identified a *cortical spectrum* of weakly laminated areas on one end and sharply laminated areas on the other end broadly across the cortex, but within lobes as well – in which the degree of lamination contributes to cortical *types*. While their results were based on quantitative assessments of histology and immunochemistry, as well as MR data in non-human primates, our results identify that transcriptomics play a role in this potential *cortical spectrum* and that the CT/M ratio serves as a useful metric in relating the balance between continuous gradients and discrete regions. For example, John and colleagues (2022) write: “Cortical types must be understood as discretizations of these continuous gradients.” We propose that the CT/M ratio serves as a metric bridging the gap between continuous gradients and discretized parcellations.

Second and third, respectively, these findings also complement recent studies reporting *transcriptomic distinctiveness* (Wagstyl et al., 2022) with visual cortex on one end and frontal cortex on the other in addition to a genetic variation along a *sensorimotor-association (S-A) cortical axis* (Sydnor et al., 2021), which has also been associated with the distribution of transmitter receptors (Goulas et al., 2021). Many terms aside – from *crazy-pavement* classically to *cortical spectrum* and *transcriptomic distinctiveness* modernly – altogether the present findings incorporate transcriptomics and a novel metric (CT/M ratio) as two features that can guide future studies striving to build mechanistic models to better understand the relationship among gradients, areal parcellations, and cortico-cortical connectomics, as well as the role of transcriptomics and the CT/M ratio as metrics mediating this complex relationship of brain structure, function, cortical networks, and genetic expression.

### Transcriptomics as a bridge connecting areas of functionally distinct cortical networks

While previous findings show that transcriptomics contribute to the arealization of a cortical network specialized for visual processing (Gomez et al., 2019), the present findings also show that areas located in different functional networks can also be more transcriptomically similar to one another than areas located in the same functional network. Specifically, even though visual areas also cluster together in the present study (lower left in Figure 3*B*), a subset of visual areas are also transcriptomically similar to areas in somatomotor cortices (3a,3b,4a,4b), secondary somatosensory cortex (OP1), and posterior parietal cortex (hiP3, hiP5, hiP6).

Additionally, even though areas FG1 and FG2 are cortically adjacent to one another in the posterior fusiform gyrus, FG1 is more transcriptomically similar to OP1 in the parietal operculum than FG2, which is more transcriptomically similar to area 5M in medial parietal cortex and PGp in lateral parietal cortex. Finally, even though Ph1 and Ph2 are adjacent to one another in parahippocampal cortex, the data-driven approach implemented here situates them in distinct transcriptomic clusters with different functions: Ph1 is transcriptomically similar to areas in auditory cortex, the parietal operculum, and posterior parietal cortex, while Ph2 is transcriptomically similar to areas in orbitofrontal cortex. While there are many other examples, the combination of these findings indicate two main points. First, despite being located in different functional networks, cortical areas may also be transcriptomically similar and share a common function despite being commonly associated to one cortical network. Second, while a large portion of the transcriptomic landscape across the cortical sheet exhibits a high degree of spatial autocorrelation where nearby anatomical regions have more similar patterns of gene expression than distal regions (Burt et al., 2020), the genes identified here that exhibit the greatest degree of differential expression show that cortical distance is not the primary indicator of transcriptomic similarity (Figure 3*D*). Instead, a likely anatomical feature underlying genetic similarity between cortically distant regions in addition to those measured here is white matter connectivity – especially so given that previous studies (Arnatkeviciute et al., 2021c) have found a high degree of transcriptomic coupling between long-range connectivity hubs over local, short-range connections. Taken together, a goal for future studies will be to determine what anatomical and functional features contribute to the intriguing finding that cytoarchitectonic areas can be cortically distant from one another and assigned to different functional networks, and yet, transcriptomically similar to one another.

### Transcriptomics contribute to different scales of organization and different parcellations of the HCC

The combination of findings across studies indicate that different sets of genes contribute to different scales of functional or anatomical organization. For example, while a sparse set of genes contribute the most variance to gradients of the HCC (Burt et al., 2018) at a meso-scale, maps within a single cortical area at the fine-scale (Gomez et al., 2021), and the parcellation of the HCC into distinct cortical areas at the macro-scale (present study), largely distinct sets of genes contribute to each type of organization. Together, these findings show that transcriptomics contribute to different scales of organization of the HCC. But, what about different functional roles of different sets of genes at the same level? For example, at the macro-scale considering different parcellations of the HCC (cytoarchitectonic vs. multimodal)? In the present study, our data-driven analysis approach showed that roughly 10% (or 8-14% if choosing different sets of genes; Supplemental Figure 5) of the identified genes were different between the cytoarchitectonic and multimodal parcellation. Furthermore, enrichment analyses showed that genes specific to a cytoarchitectonic parcellation were generally more related to cell maintenance functions that likely contribute to the cellular scaffolding necessary for cytoarchitectonic features. On the other hand, genes specific to a multimodal parcellation were more associated with ion channel and transport functions, which may be related to features related to neural activation – a component contributing to the borders of the multimodal atlas (Glasser et al., 2016). In terms of limitations, a factor that could contribute to these differences is ROI size: cytoarchitectonic ROIs tend to be larger than multimodal ROIs. However, when looking at the genes that overlap between the two parcellations, both sets are heavily enriched in synaptic function, furthering the relevance of differential expression to cell-to-cell interactions. Nevertheless, an additional limitation of the present study is that the analyses were conducted in brains from healthy elderly adults. Thus, a goal of future studies is to test whether the genes identified here also contribute to the transcriptional landscape of the developing human brain.

## Conclusion

Using a data-driven approach, we tested and found that a sparse subset of genes differentially contributed to a modern cytoarchitectonic parcellation of the HCC. The expression of this sparse subset of genes was correlated with a novel metric (CT/M ratio) that bridges the gap among i) gradients at the macroscale, ii) areas at the meso-scale, and iii) cell density at the microscale. The combination of these findings not only influences future studies examining transcriptomic contributions to the novel idea of a cortical spectrum, but also developmental and evolutionary contributions to the multimodal relationship identified across spatial scales in the present study.

## Materials and Methods

### Data

1. JüBrain: https://www.fz-juelich.de/inm/inm-1/EN/Home/home_node.html
2. Allen Human Brain Atlas (AHBA): http://brain-map.org/
3. Human Connectome Project (HCP): https://www.humanconnectome.org/study/hcp-young-adult
4. Von Economo - Koskinas Atlas: http://www.dutchconnectomelab.nl/economo/

### JüBrain cytoarchitectonic atlas

The JüBrain cytoarchitectonic atlas consists of a set of cytoarchitectonic brain areas that are defined by an observer-independent analysis of laminar cell-density profiles in 10 human post-mortem brains. Each area is represented by a 3D map that describes the maximum probability with which a certain cytoarchitectonic area can be assigned to a certain macroanatomical location in the brain. Presently, 142 areas have been mapped in the present version of the atlas, v2.9, which covers over 70% of the brain. Among them, 111 areas are located across the HCC (Figure 1*A*) with 5 cortical gap maps (Supplemental Figure 1). These 111 cytoarchitectonic regions of interest (cROIs) are the focus of the present study. The parcellation is available in both the MNI305 (Colin 27) and MNI152 template space. Given that the AHBA data is also provided in the MNI152 space, the MNI152 projection of the cROIs was used.

#### Geodesic distance

In order to calculate the distance between cROIs, the JüBrain parcellation was converted to a 32,000 vertex mesh surface so that the Connectome Workbench software (Marcus et al., 2011) could be applied. This required first transforming the parcellation from the MNI152 volume space to the fsaverage surface and then, the 32k_fs_LR surface. Once the parcellation was transformed to the appropriate surface, the *cortex* function was used to calculate the geodesic distance along the surface mesh between parcels represented in millimeters (mm). For analyses described below, the pairwise distances between regions were normalized between 0 and 1 with 0 being the absolute shortest distance between any region pair and 1 being the greatest possible distance.

### Human transcriptome data and analysis

#### AHBA

The gene expression data used in the present study were obtained from the AHBA, which is a publicly available atlas of gene expression and anatomy. The AHBA employs DNA microarray analyses to map gene expression from tissue samples taken broadly across the human brain. The dataset is based upon the combination of measurements from 6 post-mortem human brains, although not all brains were sampled identically. Cortical samples were acquired using macrodissection from every brain and submitted to normalization analyses to make measurements within and between brains comparable. Each sample was associated with a 3D coordinate from the donor’s MRI brain volume and corresponding coordinate (x, y, z) in the MNI152 space. Each tissue sample was analyzed for the expression magnitude of 29,131 genes, with 93% of known genes being acquired by at least 2 probes. For more details regarding how the microarray data were normalized, please see the following documentation: http://help.brain-map.org/display/humanbrain/documentation/.

#### Gene expression preprocessing

The collection and quality control of the gene expression data has been described previously (see Online Documentation link above). Here, we discuss the preprocessing steps relevant for the present study. The raw microarray expression data for each of the six donor brains included the expression level of 29,131 genes profiled via 58,692 microarray probes. These data were preprocessed and simultaneously aligned to the JüBrain parcellation by using the abagen package (Markello et al., 2021), which is a Python toolbox that provides a reproducible pipeline processing the transcriptomic data provided by the AHBA. While the package provides several different options across the different processing steps, the default parameters were used as recommended by (Arnatkeviciute et al., 2019), which are briefly described as follows. The first step involves removing probes whose intensity value does not exceed that of background noise and, if there are multiple probes present for a single gene, selecting the probe with the highest differential stability across donors. The next step matches the tissue samples to cROIs and then normalizes expression values for each sample across genes and for each gene across samples across each donor. The scaled robust sigmoid function was used for each normalization step. Lastly, the normalized samples were aggregated into the cROIs as the “get_expression_data” function outputs the dataframe yielding the normalized expression value for each gene within each cROI, resulting in a dataframe of 111 cROIs by 15,630 genes.

#### Gene selection

Of the genes that survived quality control implemented via the abagen package, we then sought to identify genes that were most differentially expressed across the cortex. This was done by applying PCA to the cROI expression array, which resulted in a score value for each cROI provided by the first PC (Figure 3*A*). To select for genes of interest, we then selected the genes with the highest feature weight values for PC1. To further remove genes not of interest, the data were filtered to only include ‘brain-specific genes’ as identified by Burt et a., (2018). Afterwards, the top 200, roughly 1% of the original dataset, most significantly correlated genes were identified (Figure 1*C*). This then yielded the ‘transcriptomic fingerprints’ for each cROIs consisting of the normalized expression of each of these top 200 most differentially expressed (DE) genes (Figure 3*C*).

#### Examining spatial autocorrelation

To identify any trends of spatial autocorrelation in our data, we correlated gene expression profiles between all region pairs using the 15,630 genes that survived pre-processing and plotted each pairwise Pearson’s correlation as a function of normalized geodesic distance between the two regions (Supplemental Figure 3*A*). Linear regression was performed to generate a model of spatial autocorrelation in the data (*y* = -0.6723*x* + 0.3429, mean squared error (MSE) = 0.07). Then, in order to compare spatial autocorrelation in this subset of data to the top 200 DE genes, we performed the same correlation, but only using the expression of these 200 genes, which is plotted in Supplementary Figure 3*B* as a function of geodesic distance. The MSE of these data was then calculated using the same linear model previously described. As the MSE was much higher (0.29 vs. 0.07) around the autocorrelation fit when using the data from the top 200 DE genes compared to all genes indicates much greater variance in the observed correlations. This suggests that variations in gene expression (N=200) are broadly not due to similar changes in distance between regions.

#### Exploring the relationship of gene expression among cROIs

To further explore the pattern of differential gene expression across the cortex, the transcriptomic fingerprints of the cROIs were then subjugated to an agglomerative hierarchical analysis using Euclidean distances and the Ward linkage method. This clustering information was subsequently represented in a 2-D Euclidean plane via multidimensional scaling (Figure 3*B*) to visualize the genetic similarity between cROIs.

### HCP measure of T1, T2, and cortical thickness

Group average cortical maps of myelination (T1/T2) and cortical thickness were obtained from 1,096 participants included in the Human Connectome Project (Glasser et al., 2013). In each participant, T1w and T2w scans were collected using 3 Tesla (3T) MRI and then aligned to the fsLR_32k mesh surface where the average values for both myelination and cortical thickness were taken at each vertex as in our previous work (Gomez et al., 2019).

Given that the average anatomical maps were in the HCP fsLR_space, cROI MNI coordinates were transformed into the fsLR space, which was done through the use of Freesurfer (Fischl et al., 1999; Fischl, 2012) by applying a transformation to each cROI’s surface label.

After cROIs were aligned to the average HCP fsLR surface, we then extracted the mean myelination and cortical thickness value within each cROI along with the standard deviation as in our previous work (Gomez et al., 2019). We then calculated the ratio between myelination and cortical thickness values as in our previous work (Willbrand et al., 2022). Furthermore, we quantified the relationship between PC scores and this ratio in two ways: 1) using a linear regression and 2) the correlation between the cROIs PC1 score and mean anatomical value (Figure 4*A*).

### Von Economo - Koskinas measure of cell density

Histological measurements of cellular density across the cortical mantle and layer-specific measurements of cortical thickness were obtained from the digitized version of the von Economo and Koskinas atlas (von Economo and Koskinas, 1925; Scholtens et al., 2018). The atlas contains 48 distinct cortical regions with measurements of morphological information on neuronal count, neuron size, and thickness for each cortical layer and across the whole mantle. The atlas has recently been translated into a digital version that is compatible with FreeSurfer and has been made publicly available (Scholtens et al., 2018).

To obtain the Von Economo - Koskinas measurements per cROI, the two atlases were combined in the same MNI152 volume space. Then, for each morphological measurement, the average value was calculated for each cROI via the overlapping voxels from different Von Economo - Koskinas regions. This resulted in an array of 35 different measurements for each cROI: layers 1-6 and overall for cell density, cell size, overall thickness, wall thickness, and dome thickness. Here, we considered the relationship between overall cell density across layers and layer IV thickness (both measurements normalized) to the primary axis of transcriptomic variation (Figure 4*B* and *C*).

### Multimodal Parcellation

To test if the identified genes were specific to the cytoarchitectonic parcellation of the HCC or generalized to additional parcellations of the HCC, we repeated our analyses with a multimodal parcellation of the HCC (Glasser et al., 2016; Supplemental Figure 4). To do so, AHBA data were aligned to this multimodal atlas of the HCC using the same process described above for alignment to the cytoarchitectonic parcellation of the HCC. This was done using the abagen package with the same parameters resulting in a normalized expression array of 178 multimodal ROIs by 15,630 genes. The same analyses (*gene selection* and *exploring the relationship of gene expression among ROIs*) were then performed on the multimodal expression array.

### Gene ontology analyses

To further elucidate the functional characteristics of the different, as well as overlapping, sets of genes identified from our analyses examining the relationship between i) transcriptomics and a cytoarchitectonic parcellation of the HCC and ii) transcriptomics and a multimodal parcellation of the HCC, we performed three different enrichment analyses using the following open-sourced program: ToppFun within the ToppGene Suite (Chen et al., 2009); https://toppgene.cchmc.org/enrichment.jsp. This open-sourced program detects if any functionally related genes exist within the identified set significantly higher than what would be expected by random chance (with significance values corrected using a Benjamini-Hochberg False Discovery Rate procedure). For determining if a particular gene group was significant, we chose a *p* value method of probability density function, and restricted our gene ontology identification to the following three categories: (i) GO: Molecular Function, (ii) GO: Biological Process, (iii) GO: Cellular Component. Importantly, choosing a different *p* value method (cumulative distribution function), did not significantly change the results. The first enrichment analysis focused on the genes that overlapped between the cytoarchitectonic and multimodal parcellation (Figure 5*A*), while the second and third enrichment analyses focused on genes that were unique to either the cytoarchitectonic parcellation or the multimodal parcellation (Figure 5*B*).

## Acknowledgements

Research for this project was supported by start-up funds from the University of California, Berkeley. We thank Jesse Gomez and Zonglei Zhen for important methodological guidance and discussions when setting up the initial analysis pipeline for the present study.

