## Supplemental Figures for "Transcriptomic contributions to a modern cytoarchitectonic parcellation of the human cerebral cortex"

### Supplemental Materials

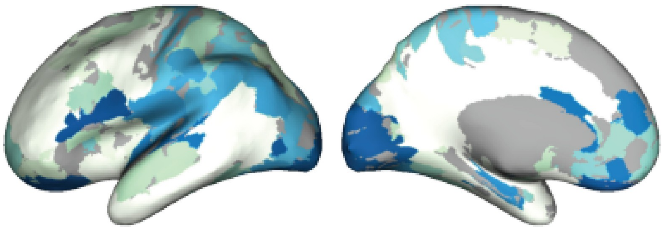

**Supplementary Figure 1. A modern cytoarchitectonic parcellation of the HCC.** cROIs in shades of blue and *gap maps* in white from Amunts et al., 2020 in the MNI 152 stereotaxic space.

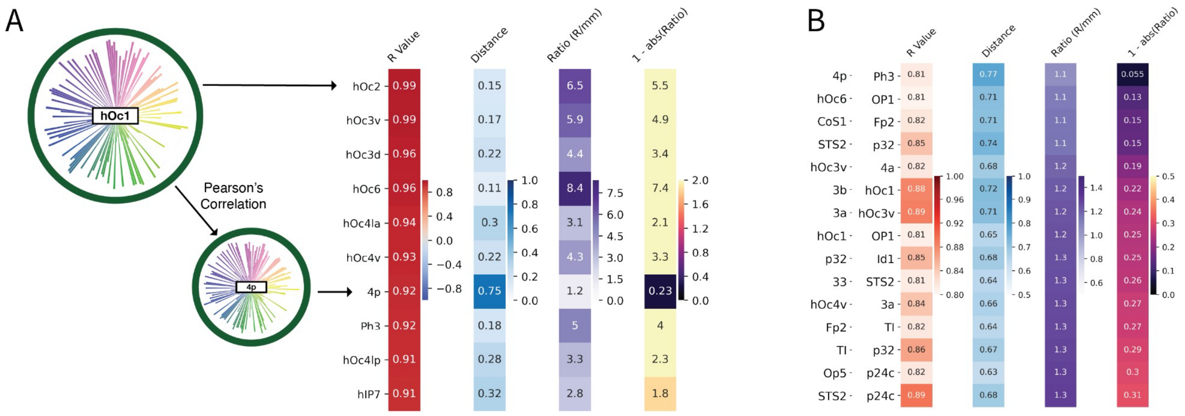


**Supplementary Figure 2. Identification of region pairs that are spatially distant, but highly correlated in gene expression.** (A) A schematic representation of how the transformed ratio value shown in the column on the far right is calculated using hOc1 as an example. The first column represents Pearson's correlation coefficient in the expression of the top 200 DE genes shown in the two example fingerprints on the left with hOc1 as the base region and area 4p as one of the comparison regions. The top 10 regions with the highest correlation and their values are displayed, ranging from hOc2 to hIP7. The second column shows the normalized geodesic distance value between hOc1 and the top 10 correlated regions. The third column is simply the ratio of the correlation value divided by the normalized distance value. Finally, the 4th column represents the 'transformed ratio' values that are used to quantitatively identify region pairs that are highly correlated in their expression of the top 200 DE genes, but spatially distant from one another. The transformed ratio value was calculated by taking the absolute value of the ratio value and subtracting it from 1 so that any region pair with a transformed ratio value close to 0 will represent region pairs that are not only highly correlated, but also distal from one another. For example, the region that has the lowest transformed ratio value with hOc1 is area 4p (R = 0.92 and normalized distance = 0.75). (B) The top 15 region pairs with the lowest transformed ratio value filtered to R values > 0.8. The 4 columns are the same columns described in (A), but ordered by the transformed ratio value with area 4p in the somatosensory cortex and Ph3 in the parahippocampal gyrus showing the lowest transformed ratio (R = 0.81 and normalized distance = 0.77).

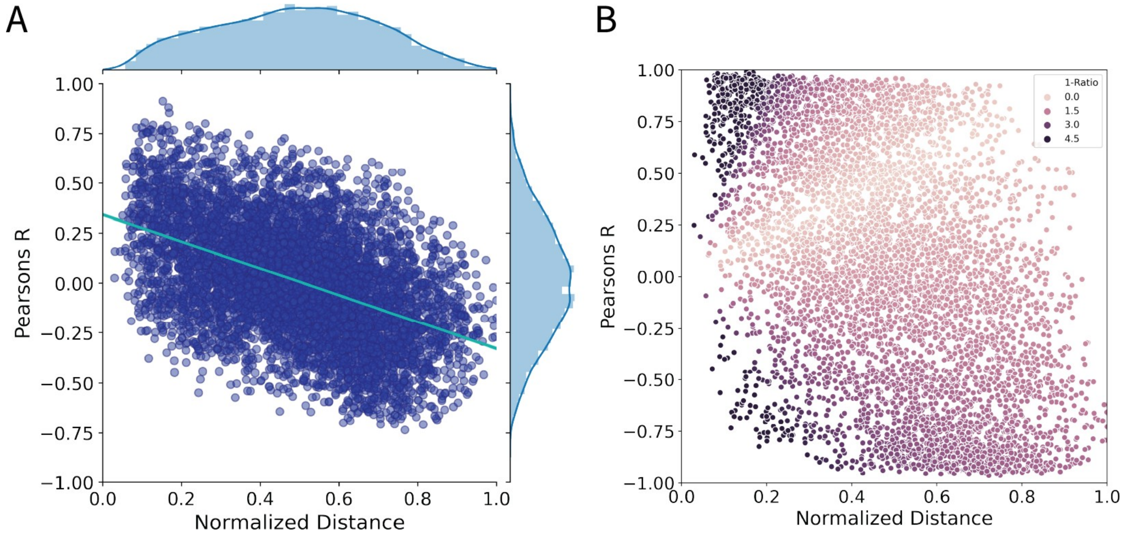


**Supplementary Figure 3. Spatial autocorrelation of transcriptomic profiles between region pairs.** (A) Correlation in gene expression of the 15,630 genes surveyed post-processing between all pairs of regions ($111^2$ region pairs total). Correlation is represented by Pearson's R value on the y-axis with normalized geodesic distance (Materials and Methods for more details) between two regions being represented on the x-axis. Each blue point corresponds to a region pair's correlation in gene expression as a function of distance. Distribution of correlation values is shown on the right y-axis and distribution of normalized geodesic distance values is shown on the top x-axis. Linear fit through the data is plotted in cyan ($y = -0.6723x + 0.3429$, mean squared error (MSE) = 0.07). (B). Similar to (A), correlation in gene expression between region pairs is plotted as a function of normalized geodesic distance. However, correlation is calculated based on the expression of the top 200 differentially expressed (DE) genes. In other words, correlation between the transcriptomic profiles or 'fingerprints' of regions as illustrated in

Figure 3*C*. The hue of the datapoints represents the transformed ratio value calculated (see Supplemental Figure 2*A* for details) between each pair of regions. To quantify differences in autocorrelation when examining the gene expression of all 15,630 genes vs. the top DE 200 genes, the MSE was calculated for the data shown in *B* using the linear model shown in *A* resulting in an MSE of .29. Thus, the data shown in *B* exhibits much greater variance around the linear model of spatial autocorrelation generated, indicating that the 200 genes analyzed are less prone to trends in spatial autocorrelation.

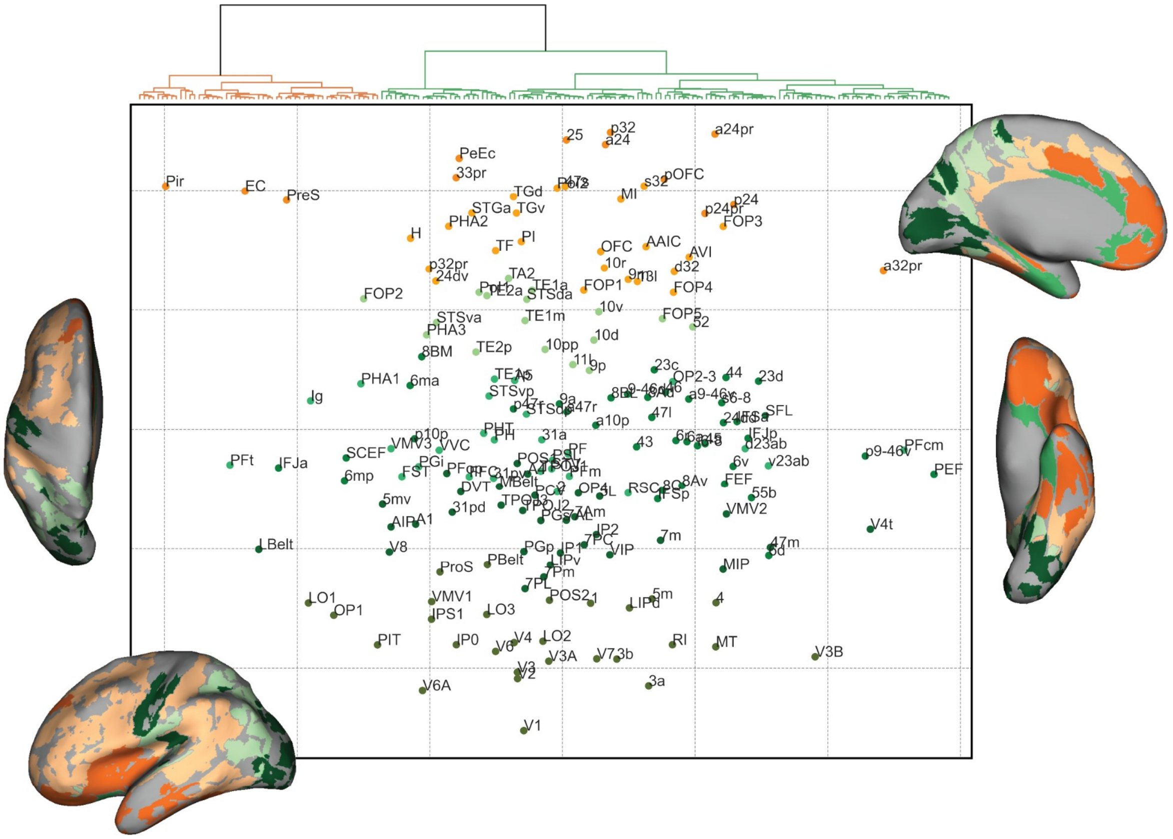


**Supplementary Figure 4. Top differential gene clustering of multimodal ROIs from Glasser and colleagues (2016).** Differential gene expression, clustering, and MDS analyses were also performed for AHBA data aligned to the Glasser Multimodal Parcellation resulting in a replication of Figure 3*B*, but with multimodal-aligned transcriptomic data. Similarly to the cytoarchitectonic parcellation, there are two clusters at the highest level separating primary sensory regions from frontal and insular regions with similar subclusters shown in the lighter shades of green and orange. Also similar results occur for the MDS analyses:  V1 is at one end of the MDS axis with insular regions at the other end. Color coded clusters are also depicted topographically on the MNI-152 inflated surfaces shown on the left and right, respectively.

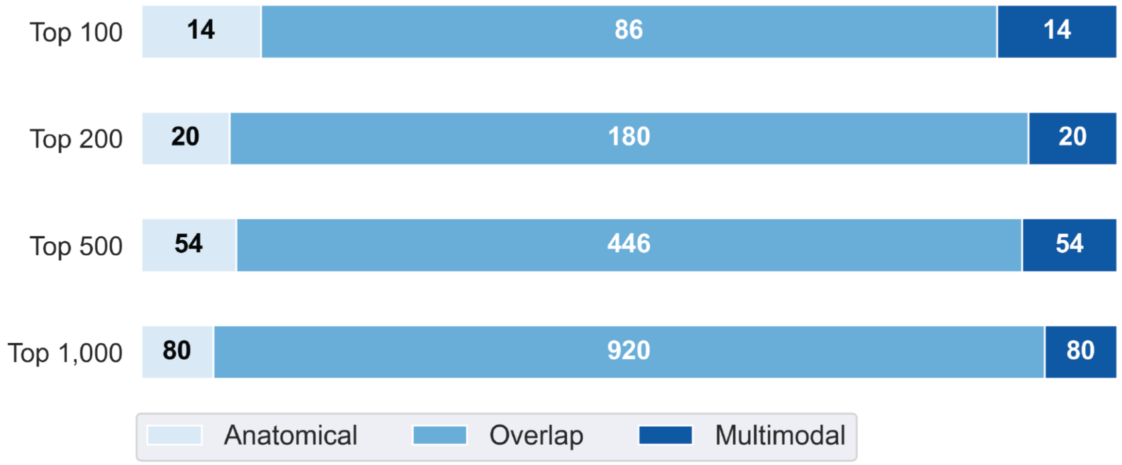


**Supplementary Figure 5. Top differential gene comparison between parcellations.** Venn-diagrams represented as horizontal bars for the overlapping genes between the two different parcellations ('Anatomical' being the cytoarchitectonic atlas). Different rows correspond to the different sized N of top differential genes (N = 100, 200, 500, 1,000) that were 'brain-specific' (see Burt et al., 2018). For the top 200 differentially expressed genes, 180 of the 200 genes overlap and the other 20 genes are unique to each parcellation. Through each iteration of N genes, at least 8% of genes are unique to either parcellation with the most being 14%.

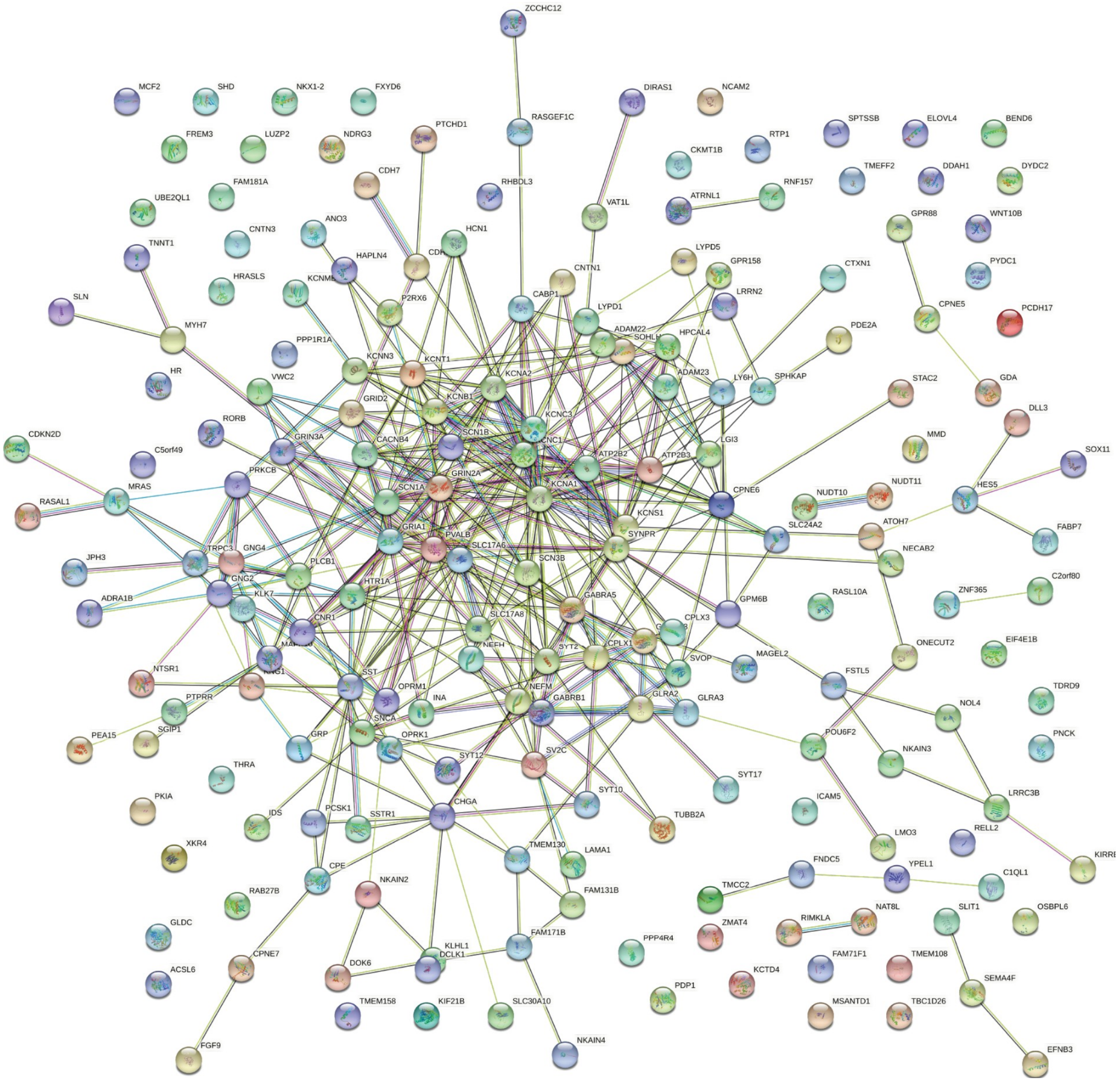


**Supplementary Figure 6. Network graph of gene enrichment connections.** In addition to enrichment analyses performed using the GO database (see Figure 5), a protein-protein interaction graph was generated using STRING. Each connection between each gene has a score that represents the degree of confidence of a true interaction between the corresponding proteins for those genes and different colored lines represent different types of interactions. In sum, the strongest interactions are between the ion-channel function genes (i.e. KCNA and SCN gene families), neurotransmitter genes (CNR1, GRIN2A, etc.) and cell maintenance genes. Other interactions include transcription factors (SOX11 and HES5) and early developmental genes (SLIT1 and CNTN gene family).
